# Duck pan-genome reveals two transposon-derived structural variations caused bodyweight enlarging and white plumage phenotype formation during evolution

**DOI:** 10.1101/2023.01.28.526061

**Authors:** Kejun Wang, Guoying Hua, Jingyi Li, Yu Yang, Chenxi Zhang, Lan Yang, Xiaoyu Hu, Armin Scheben, Yanan Wu, Ping Gong, Shuangjie Zhang, Yanfeng Fan, Tao Zeng, Lizhi Lu, Yanzhang Gong, Ruirui Jiang, Guirong Sun, Yadong Tian, Xiangtao Kang, Haifei Hu, Wenting Li

## Abstract

Structural variations (SVs) are a major source of domestication and improvement traits, however SV profiles of duck and their phenotypic impacts largely hidden. We present the first duck pan-genome constructed using five genome assemblies capturing ∼40.98 Mb new sequences. This pan-genome together with high-depth sequencing data (∼46.5X) identified 101,041 SVs, of which substantial proportions were derived from transposable element (TE) activity. Many TE-derived SVs anchoring in a gene body or regulatory region are linked to duck’s domestication and improvement. By combining quantitative genetics with molecular experiments, we dissect how TE-derived SVs change gene expression of *IGF2BP1* and generate a novel transcript of *MITF*, shaping bodyweight and white plumage. In the *IGF2BP1* locus, the TE-derived SV explains the largest effect on bodyweight among avian species (27.61% of phenotypic variation). Our findings highlight the importance of using a pan-genome as a reference in genomics studies and explore the roles of TE-derived SVs in trait formation and in livestock breeding.

## Introduction

Phenotypic variation in livestock is shaped by genetic variation accumulated from wild ancestors to indigenous breeds and subsequent breeding. A major goal of animal genetics and breeding is to identify and dissect how functional genes or variants diversify phenotypes. Many phenotypes in livestock are affected by structural variations (SVs), especially in poultry, including feathered legs, crest, and body size of chicken [1-3], as well as plumage pigmentation of pigeon [4]. Previous studies suggest that SV generation is associated with the transposable element (TE) mediated imprecise duplication, insertion, deletion, and reshuffling of host genome sequences during genome evolution [5-8]. In addition, TE-derived SVs contribute to phenotypic variations in vertebrates by affecting the transcriptional regulation via donating promoter or enhancer sequences, modifying 3D chromatin architecture that regulates the host genes’ expressions and leading to *de novo* gene birth [6, 9, 10]. Examples include henny feathering in chicken [11], white coat color in buffalo [12], secondary palate development [13], and embryonic implantation in humans [14]. However, a reference genome derived from one individual cannot fully capture a species’ genetic diversity, leaving the majority of SVs poorly resolved and their phenotypic impacts largely hidden.

A pan-genome, which captures the complete genetic variations of a species, enables an abroad survey of SV landscapes using population-scale sequencing data, resulting in better characterizations of SVs and understandings of their impacts on phenotypic variations. Duck, in addition to its agricultural and economic importance, has extensive genetic variation resources which descend from wild mallard (*Anas platyrhynchos*) and spot-billed duck (*Anas zonorhyncha*) [15-17] and phenotypic variations in their morphology, productivity, and behaviour [18], making it a potential model to investigate the significance of SVs in animal biology. Here, we constructed the first duck pan-genome using five genome assemblies and investigated SVs in 12 populations of 131 ducks (wild, native, and commercial breeds) with high-depth sequencing data. By identifying the associations between SVs and TEs across the pan-genome, we found an increase in the occurrence probability of SVs linked with TE activity. To demonstrate the significance of these TE-derived SVs, we linked these SVs with domestication and improvement traits. We found these SVs affected the expression level and contributed to new transcript generation of the causative genes regulating the quantitative bodyweight trait and qualitative plumage pattern. Our work expands the understanding of the importance of the pan-genome and underlines the pronounced effect of TE-derived SVs on phenotype formation.

## Results

### Construction of duck pan-genome

We constructed the first duck pan-genome using a combination approach of *Psvcp* [19] and *PPsPCP* [20] pipelines. Five published duck genomes were used to construct the duck pan-genome, which consists of three Pekin duck genomes (commercial breed), one Shaoxing duck genome (indigenous breed), and mallard duck genome (wild relatives) (Additional file 1: Table S1). The approach we used for constructing the duck pan-genome is visualized in Fig. 1a. Briefly, duckbase.refseq.v4 genome was aligned to the initial reference genome (ZJU1.0), then insertions longer than 50 bp were identified and placed in the ZJU1.0. This process was further iterated by CAU_Pekin2.0, CAU_Laying_1.0, and ASM8764695v1 in order of increasing phylogenetic distance to ZJU1.0 (Additional file 2: Fig. S1a), and thus the Pan-genome.1 was generated. Subsequently, four query genomes were aligned to Pan-genome.1, respectively, while novel contigs longer than 500 bp were retained after removing redundancy. Novel contigs and Pan-genome.1 were merged into the final duck pan-genome (Additional file 1: Table S2). The duck pan-genome identified ∼ 40.98 Mb additional sequences that were absent from the reference genome (ZJU1.0), encoding 329 high-confidence genes with intact coding regions. In the *Psvcp* pipeline, 21,403 insertions were identified and placed into 30 chromosomes, cumulatively encoding the genomic length of ∼7.42 Mb (Fig. 1b). Novel contigs generated from the *PPsPCP* pipeline comprised 1,830 sequences cumulatively encoding genomic sequence length of ∼33.56 Mb (Fig. 1c).

**Fig. 1.**
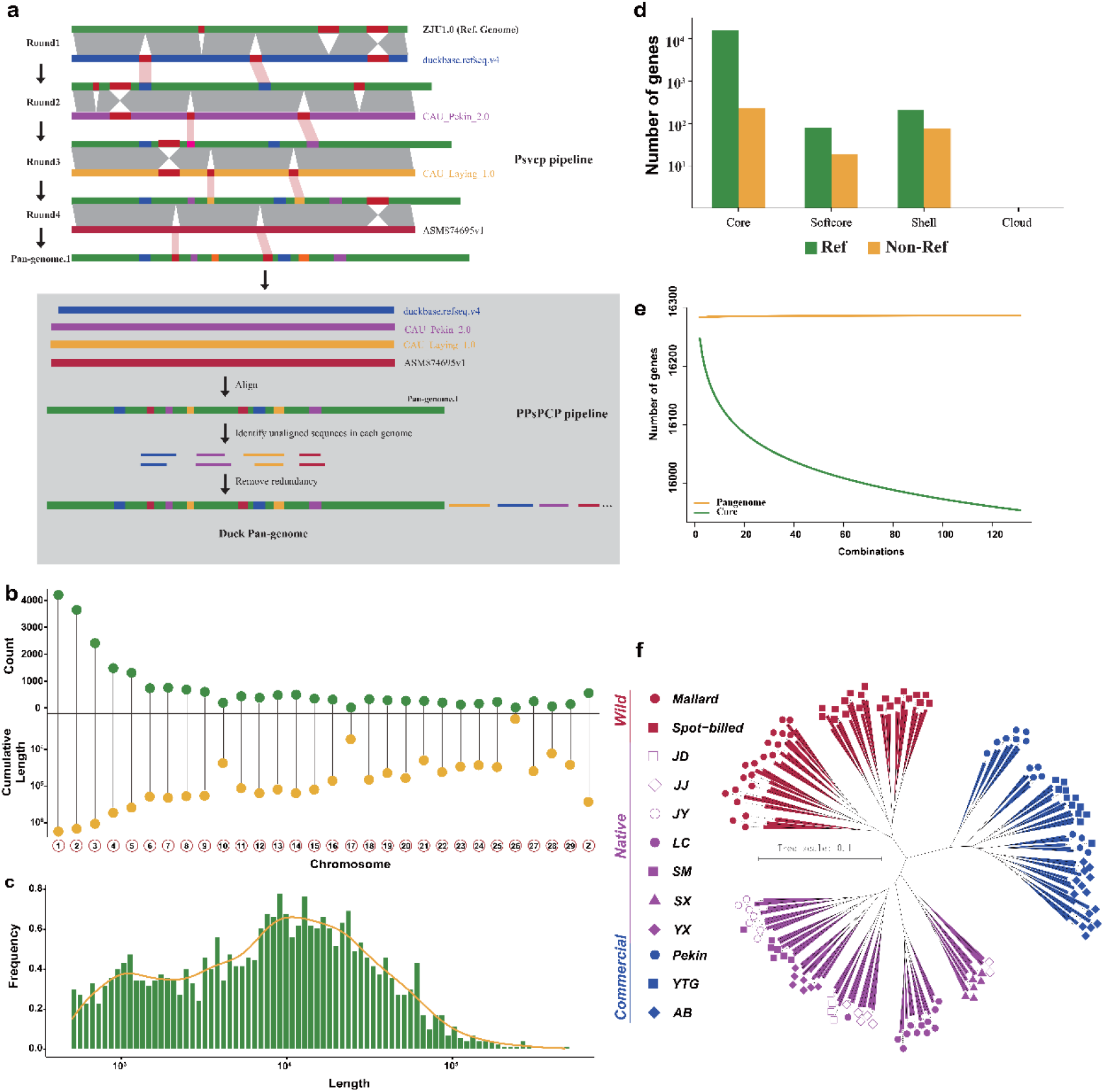
Pan-genome of Duck. (a) Schematic of duck pan-genome construction. (b) Statistics of newly identified sequences placed in chromosomes via Psvcp pipeline. (c) Length distribution of novel sequences identified via PPsPCP pipeline. (d) Classification of Pan-genome genes. (e) Pan-genome modeling. The model describes the trend of core gene number and pan-genome number with each genome added. This model reaches a plateau when additional genomes are added, indicating a closed pan-genome. (f) Phylogenetic tree constructed based on SVs with GTR model. Wild: wild group, including mallard and spot-billed (Chinese spot-billed duck); Native: indigenous duck breeds, including JD (Jinding duck), JJ (Jingjiang sheldrake), JY (Jinyun sheldrake), LC (Liancheng white), SM (Shan sheldrake), SX (Shaoxing), and YX (Youxian sheldrake) duck; Commercial: commercial duck breeds, including Pekin, YTG (Cherry Valley), and AB (Grimaud freres) duck.

We further generated 131 high-depth sequencing data from 3 commercial duck breeds, 7 indigenous breeds, and 2 wild species, with an estimated average depth of up to 46.5× through the pan-genome (Additional file 1: Table S3). We categorized genes in the duck pan-genome according to their gene PAV (Presence/Absence variation) frequencies in all duck breeds. A total of 15,906 (97.67%) core genes were shared by 131 individuals. The remaining 380 (2.33%) dispensable genes comprised 98 softcore, 282 shell, and 0 cloud genes, which were defined with a present frequency of more than 99%, 1-99%, and less than 1% respectively (Fig. 1d). This duck pan-genome showed a higher proportion of core gene content compared to that of human (96.88%) [21], chicken (76.32%) [1], and mussel (69.2%) [22]. Gene Ontology (GO) analysis revealed that dispensable genes were enriched in terms including immune response, sensory perception of smell, and G protein-coupled receptor signaling pathway (Additional file 2: Fig. S2). Evidence from pan-genome modelling revealed a closed pan-genome with an estimated total of 15,959 genes (genes on sex chromosomes were excluded, Fig. 1e). This suggests the current duck pan-genome assembled included nearly complete genetic diversity.

### Change of SV frequency during duck domestication and improvement

Four different SV detection tools LUMPY [23], Delly [24], GRIDSS [25], and Manta [26] commonly used for SV detection in animals [27] and plants [28] were employed. SVs were identified based on our pan-genome using high-depth sequencing data (average 46.5×) with support by at least two SV detection tools to reduce the false positive SV discovery. As a result, we generated a final set of 101,041 SVs among 131 duck genomes. Due to the limitations of short read sequencing and its SV detection algorithms, it may lead to biases in SV discovery [29]. To estimate the accuracy of SV detection in this study, we validated the consistency of four randomly selected SVs detection at the population level using the PCR genotyping method. Based on the PCR results of 131 samples × 4 long fragment SVs genotypes, the average accuracy rate is 87.4% (Additional file 1: Table S4), which is similar to the 88% SV calling accuracy reported by a peach SV discovery study [28]. To demonstrate the advantage of using a pan-genome in SV detection, we found that using the pan-genome as the reference can lead to the identification of an additional 27,533 autosomal SVs than using the single reference genome ZJU1.0.

SV-based genetic analysis revealed that 131 individuals clustered into three major groups: wild, native, and commercial, as shown in the phylogenetic tree, principal component analysis and population structure (Fig. 1f, Additional file 2: Fig. S3). Evolutionary relationships inferred from SVs are consistent with the evidence inferred from SNPs (Additional file 2: Fig. S4). To uncover the changes in SVs occurrence frequencies during duck domestication and subsequent breeding improvement, we conducted two sets of comparisons between the wild and native breeds for domestication (Fig. 2a, c) and between the native and commercial breeds for breeding improvement (Fig. 2b, d). The overlap of significant results between a Fisher’s exact test [1, 30] and the Fixation index (F_ST_) value [31] was defined as the SVs selected during domestication or improvement. We observed occurrence frequencies of 999 SVs showing significant differences between native ducks and wild ducks, with 382 SVs increased and 617 SVs decreased in frequencies (Fig. 2e-f, and Additional file 1: Table S5). GO analysis indicates that genes adjacent to SVs selected regions (potentially selected genes) during domestication were enriched in functions associated with neuron development, anatomical structure morphogenesis, cell morphogenesis, response to bacterium process, etc. (Fig. 2g-h). Due to captivity and selection during domestication [32], a 24%-35% reduction in brain size was reported in ducks [33, 34], leading to neuron development alteration of visual and trigeminal systems in the telencephalon. Potentially selected genes enriched in structure morphogenesis and cell morphogenesis may contribute to the increase in body size of native ducks during domestication [35], which is similar to the observation during yak domestication [36]. Additionally, potentially selected genes involved in response to bacteria may result from the living environment alteration and changes in pathogen pressure during domestication. Besides, 518 SVs increased and 435 SVs decreased in frequencies were detected between commercial ducks and native ducks (Fig. 2e-f, and Additional file 1: Table S6). Genes affected by these selected SV during improvement were enriched in cell adhesion, reproduction, spermatogenesis, peptidyl-tyrosine dephosphorylation (Fig. 2i). The fact that reproduction associated genes were under further selection during breeding improvement, is in line with higher performance of egg production in commercial breeds compared with native ducks. Intriguingly, we note that two well-studied genes associated with productive and morphological traits, respectively, were adjacent to deletions with significant frequency changes during improvement (Fig. 2b). A ∼7.0 kb deletion (ID: DEL00154411) located at the upstream region of *IGF2BP1* shows a higher occurrence frequency (0.89) in native ducks compared with an extremely lower frequency (0.012) in commercial ducks (FDR-adjusted p = 5.54E-34 and F_ST_ = 0.86). Another ∼6.6 kb deletion (ID: DEL00130156) located in an intron of *MITF* presents a significantly higher frequency (0.93) in native ducks while completely absent in commercial ducks (FDR-adjusted p = 4.55E-39 and F_ST_ = 0.92).

**Fig. 2.**
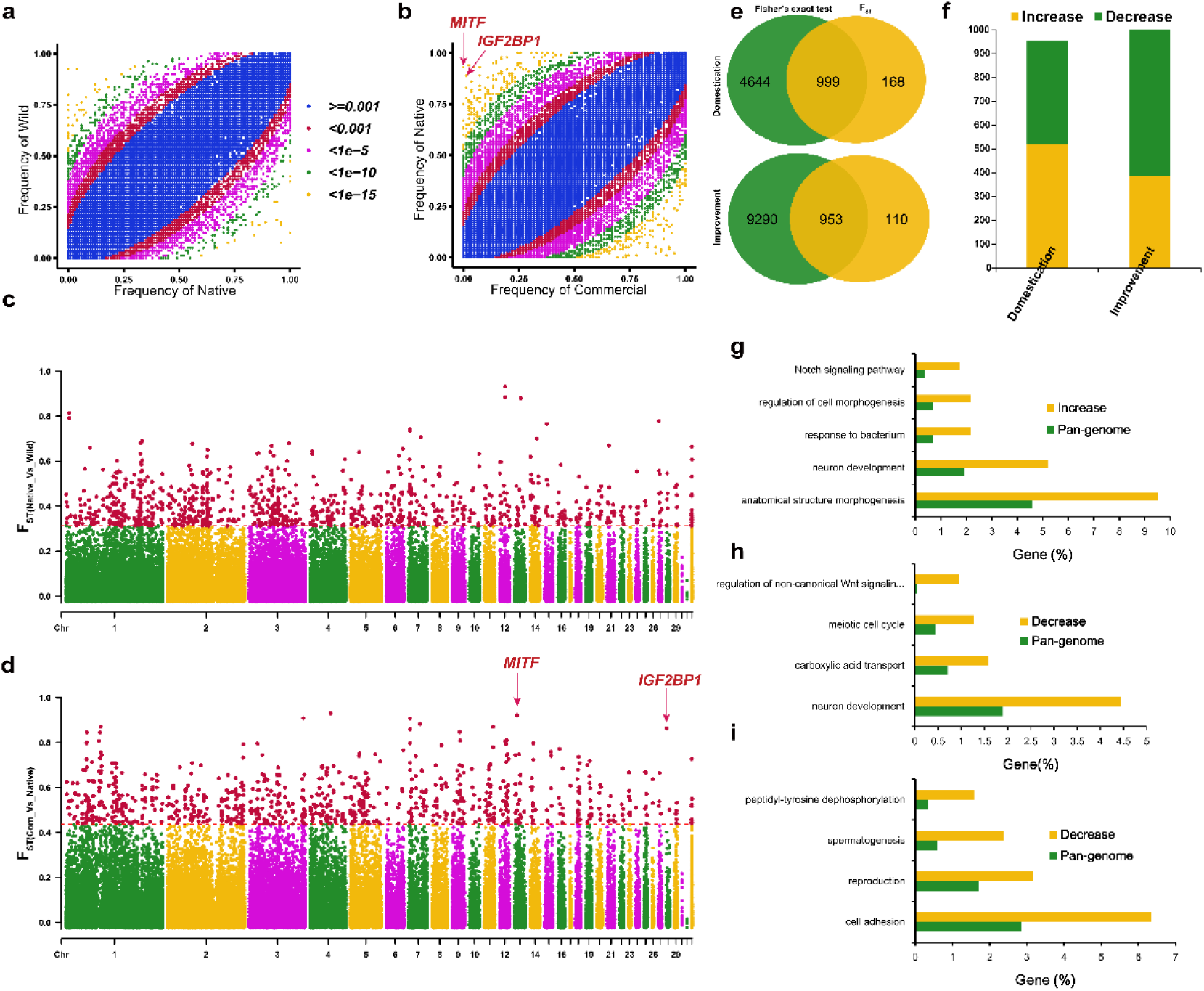
Selection of SVs during duck domestication and improvement. Scatter plots showing SV occurrence frequencies in (a) Wild and Native (comparisons for domestication) and in (b) Native and Commercial groups (comparisons for improvement). Manhattan plots showing F_ST_ between (c) Native and Wild groups, and between (d) Native and Commercial groups. Top 1% was defined as significant SVs. (e) Venn plot showing the overlap between Fisher’s exact and F_ST_ significant SVs. (f) Classification of significant SVs during evolution. Enriched GO terms in genes closest to (g) frequency increased and (h) decreased SVs during domestication, and (i) decreased SVs during improvement.

### Transposon-derived structural variations are linked to duck domestication and improvement

Using de novo-prediction and sequence similarity detection methods, around 24.37% of the newly identified sequences were annotated as TEs, while only 9.49% of entire duck pan-genome sequences were TEs. Accumulative evidence revealed that TEs offer the fodder for pan-genome dynamics resulting from its virtue of replication and mobilization [37, 38]. TEs transposition propagation experienced insertion and removal processes that easily lead to SV generation because of imprecise manipulation, reshuffling of host genome sequences, and recombining of highly homologous regions [6-8]. These factors implied the possibility of SV generation correlated to TE distribution during genome evolution. Therefore, we further investigated the classification, abundance, and length of TEs across the pan-genome (Additional file 2: Fig. S5 and Fig. 3a) and estimated the correlation between TEs and SVs using a sliding window approach with different window sizes (Fig. 3b). Significant presence correlations between SVs and TEs were observed for all the tested window sizes (Fig. 3b). The presence frequency of SVs significantly increased when TEs were present. In addition, 59.2% of identified SVs larger than 100 bp match at least one TE, of which 58.7% show at least half of the sequence overlap with TE (high TE-derived SVs) (Fig. 3c). Of these, 10.48% and 10.79% of high TE-derived SVs were located within a gene body and 5 kb of the upstream region of a gene (Fig. 3d).

**Fig. 3.**
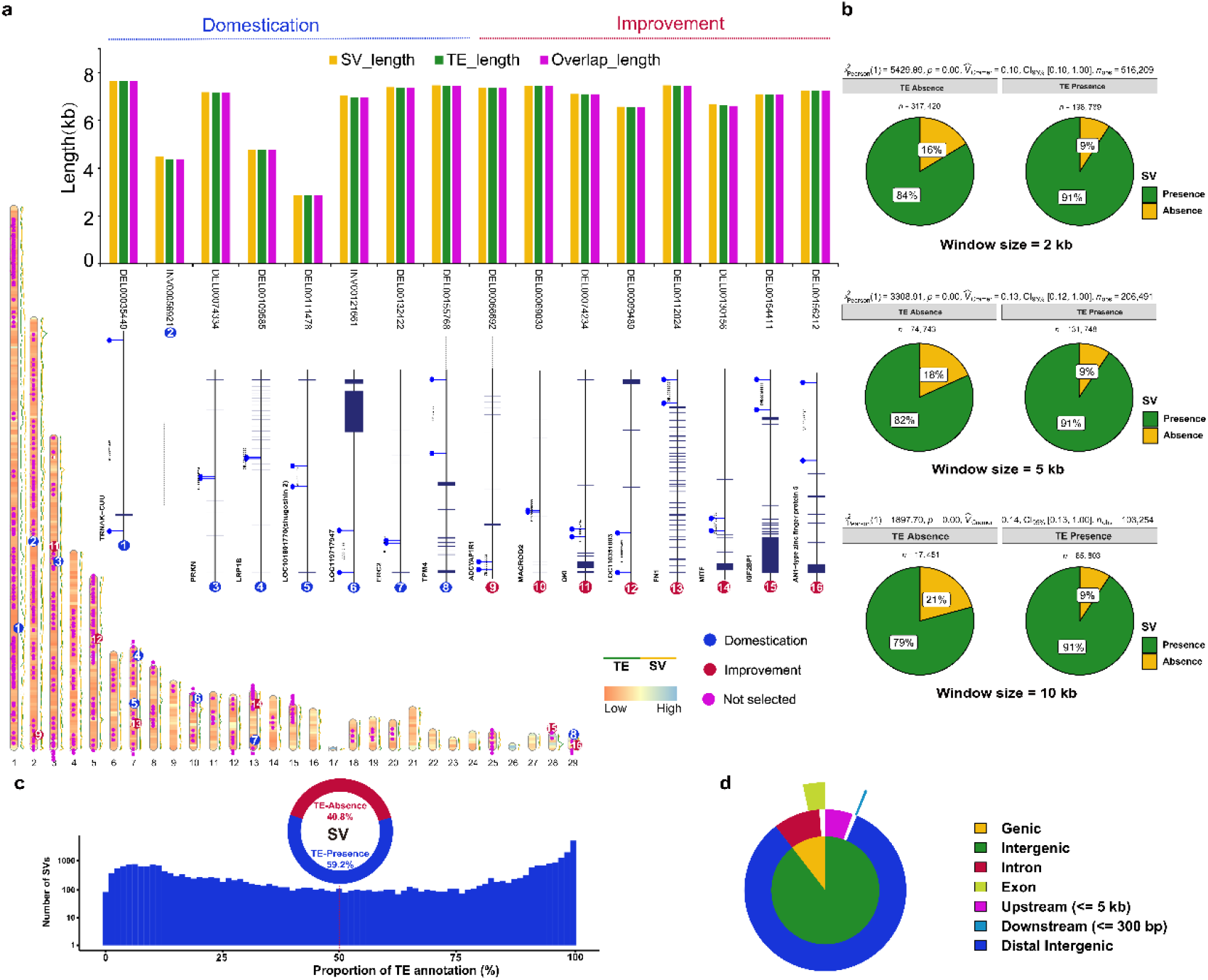
Transposable elements (TEs) annotation and their association with SVs based on duck pan-genome. (a) Localizations of SVs and TEs across the duck pan-genome. Alone with the schematic chromosomes, the green line represents the TE count of each Mb window; the yellow line represents the SV count of each Mb window; the purple dots above the schematic chromosomes show the position of intact-TE derived SVs, while red and blue dots represent the selected SVs during domestication and improvement, respectively. Imbedded histogram shows the length of SV, TE and the overlap length between intact-TE and matched SVs under selection; Positional relationships between fifteen intact-TE derived SVs and their nearest genes were labeled below, which SVs located between two blue sticks. (b) Co-occurrence probability between SVs and TEs across the pan-genome investigated by the Chi-square tests. (c) The proportions of SVs matched to TEs and the distribution of overlap proportions with TE (below). (d) Percentage of genomic features with high TE-derived SVs.

To reveal which SVs were entirely derived from intact TEs insertion across the duck pan-genome, 5,240 intact TEs were identified using the Extensive De-novo TE Annotator (EDTA) pipeline, and 392 SVs were matched to identified TEs with sequence overlap of more than 95% (Additional file 1: Table S7). Of these intact TE-derived SVs, we found that the occurrence frequencies of 16 SVs were significantly altered during the evolution of domesticated ducks, including 8 SVs during domestication and 8 SVs during improvement (Fig. 3a). Of these 16 SVs, 8 located within the intron of genes and 7 were found in the upstream or downstream (from 0.88 kb to 492.5 kb) of genes (Fig. 3a). Interestingly, these intact TE-derived SVs included the two involving well-studied genes, as mentioned above: DEL00154411 locates upstream of *IGF2BP1* and DEL00130156 within the intron of *MITF*. SVs triggered by TEs could alter the expressions of the nearest genes or generate novel transcripts of host genes ^14^, which is likely to diversify the phenotypes, including quantitative and qualitative traits. Subsequently, we took the SVs in *IGF2BP1* and *MITF* as examples to decipher how TE-derived SVs can drive duck phenotypic evolution.

### Insertion of a 6,945 bp Gypsy transposon into the promoter region of *IGF2BP1* enlarges duck bodyweight via increasing *IGF2BP1* expression

*IGF2BP1* was reported to be associated with the bodyweight of duck, in which a higher level of mRNA is correlated to a higher bodyweight [35]; however, the causative variant of *IGF2BP1* responsible for bodyweight variability remains unresolved. Our pan-genome analysis and subsequent PCR Sanger-sequencing revealed a 6,945 bp genome sequence insertion in the *IGF2BP1* promoter region, the presence and absence of which define H (Heavy) and L (Light) alleles of the gene, respectively (Fig. 4a-b, and Additional file 2: Fig. S6a). The result from allele-specific PCR demonstrated that the H allele is nearly fixed in the commercial breeds, consistent with their higher body weight comparing to the native and wild breeds [35]. In contrast, the L allele is dominant in the native and wild breeds (Fig. 4c-d, and Additional file 2: Fig. S6b), consistent with the increase in the frequency of this SV (H allele) during improvement (Fig. 2). Considering that the 6,945 bp insertion could be a linked marker of the causal variant, we further investigated the genetic polymorphisms flanking the 6,945 bp inserted sequence. We calculated the absolute allelic frequency differences (absAFdif) between the individuals from the commercial duck breeds with higher body weights and the ducks with lower body weights. Among the 90 investigated genetic variants, the 6,945 bp insertion shows the highest absAFdif (0.90), indicating that this insertion which located in the promoter region of *IGF2BP1* is the top candidate variant responsible for the heavier body weight in duck.

**Fig. 4.**
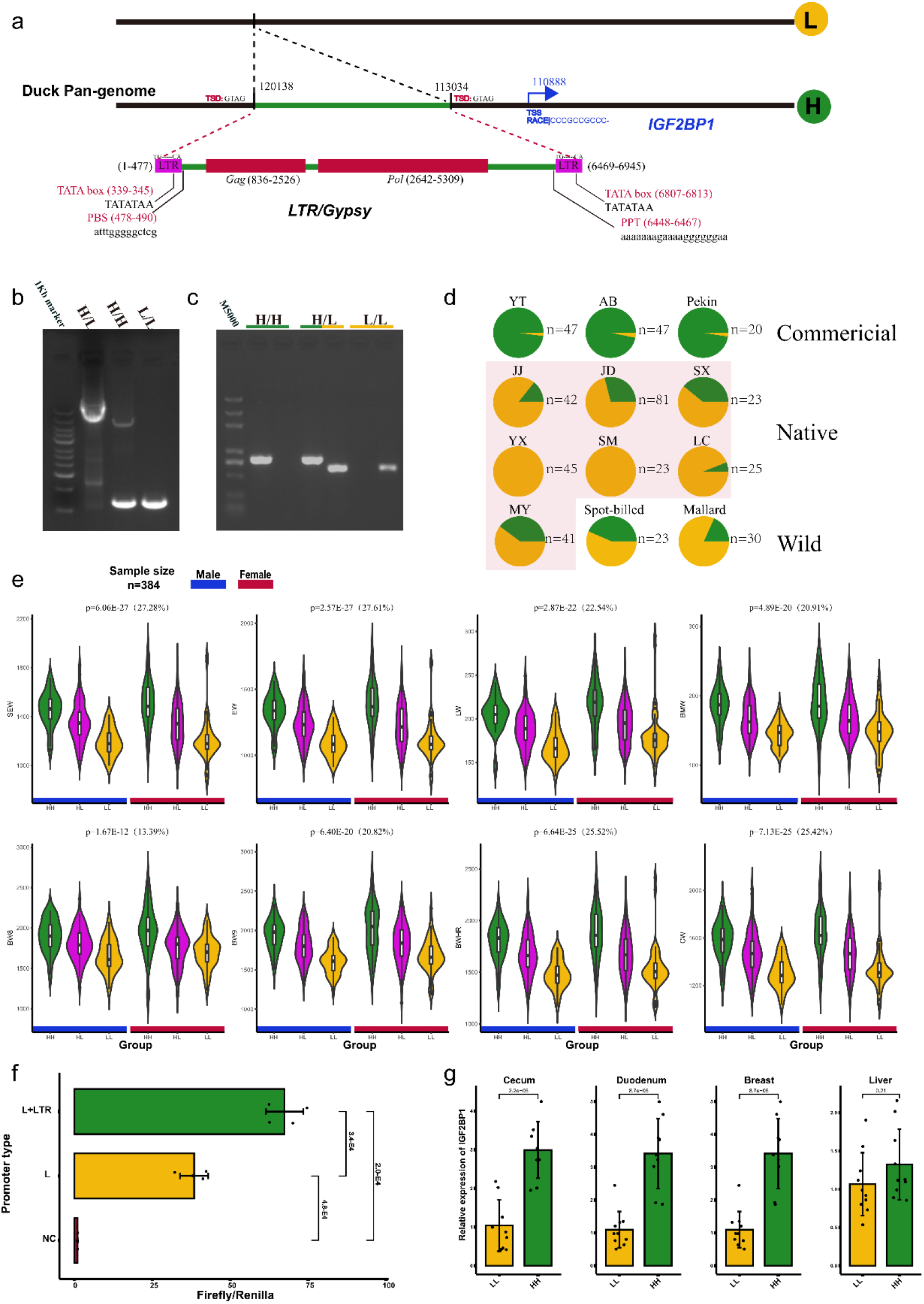
The Gypsy element regulated *IGF2BP1* expression and was associated with bodyweight traits. (a) The structure of the Gypsy element is anchored in the *IGF2BP1* promoter region. (b) Gel plots for the PCR genotyping and (c) allelic-specific PCR genotyping. (d) Allelic frequencies of the *IGF2BP1* promoter insertion in the validated populations by allelic-specific PCR genotyping. (e) Single-marker genotype association of *IGF2BP1* promoter insertion in validated duck populations. The percentage in the brackets is the proportion of phenotype variance explained by the insertion. EW, evisceration weight; SEW, semi-evisceration weight; CW, carcass weight; LW, leg weight; BMW, Breast muscle weight, BW8 and BW9, bodyweight at the age of 8 and 9th week; BWHR, bodyweight at slaughter. (f) Comparison of transcriptional activity between pGL3-L and pGL3-L+LTR recombinant plasmids representing different genotypes of *IGF2BP1* regulatory region. The significance level was analysed by a two-tailed Student’s *t*-test. (g) Comparison of mRNA expression of *IGF2BP1* between HH (YTG) and LL (SM) ducks in four tissues at 1 day of age. P-values were calculated using a two-tailed Student’s *t*-test.

Single-marker association analysis using a Liancheng white duck × Kaiya duck F2 population demonstrated that this SV is significantly associated with eight bodyweight and carcass weight traits which include evisceration weight (EW), semi-evisceration weight (SEW), leg weight (LW), breast muscle weight (BMW), bodyweight at the age of 8th and 9th weeks (BW8 and BW9), bodyweight of feather removed at slaughter (BWHR) and carcass weight (CW) (Fig. 4e). Regarding these traits, the HH genotype was always linked to a higher performance of production compared to the LL genotype. Of these, associations are most significant in EW (p = 2.57E-27) and SEW trait (p = 6.06E-27), and this locus accounts for 27.61% and 27.28% of phenotypic variations, respectively. The other traits affected by the HH genotype also presented a strong effect, with more than 20% of phenotypic variation explained for all other traits except BW8 (Fig. 4e). For BW8, HH explained 13.39% of phenotypic variation.

The 6,945 bp insertion was predicted to be an intact TE-derived SV. As a result, this TE-derived SV belongs to the Gypsy LTR superfamily with two intact LTRs as well as *gag* and *pol* coding genes (Fig. 4a). Retrotransposons can alter host gene expression or generate novel fusion transcripts [9]. After we investigated *IGF2BP1* transcription, evidence from the rapid amplification of cDNA ends (RACE) shows that no fusion transcript of *IGF2BP1* was generated and the transcription start site (TSS) was positioned at Chr28:110,888 (Duck pan-genome). To verify the effect of this TE on the transcriptional activity, a recombinant plasmid pGL3-L+LTR was defined to represent the TE insertion because of the incapacity to insert the whole TE into the pGL3 plasmid. The transcriptional activity of pGL3-L+LTR was significantly higher than that of pGL3-L (Fig. 4f). Additionally, mRNA expression of the ducks with *IGF2BP1* HH genotype (YTG) was significantly higher than those with the LL genotype (SM) in almost all investigated tissues at day 1 of age (Fig. 4g). RNA-seq analysis further confirmed this result, showing that the expression level of *IGF2BP1* of LL genotype (mallard) was lower than that of HH genotype (Pekin) in liver and sebum tissues at 2, 4, and 6 weeks of age (Additional file 2: Fig. S6c). Based on the above evidence, we conclude that a 6,945 bp Gypsy element inserted into the promoter region of *IGF2BP1* is likely the only functional variant and could increase the duck bodyweight by increasing *IGF2BP1* expression.

### A 6,634 bp Gypsy element inserted into the intron of *MITF* generates a chimeric transcript that contributes to the white plumage phenotype in duck

Previous studies showed that a ∼6.6 kb insertion in the intron of *MITF* was associated with the white plumage phenotype of Pekin and Cherry Valley duck [35, 39]. However, the molecular mechanism of this insertion remains unknown. Our pan-genome analysis and subsequent PCR Sanger-sequencing identified this insertion’s detailed location and sequence, which is 6,634 bp in length. The presence and absence of this insertion are defined as W (White) and C (Colored) alleles, respectively (Fig. 5a). Results from the allele-specific PCR demonstrate that the WW genotype is completely absent in ducks with colored plumage while completely fixed in commercial white plumage ducks (Pekin, YTG, and AB) (Fig. 5b, and Additional file 2: Fig. S7a-d). WW homozygote is linked to the white plumage phenotype, while WC heterozygote and CC homozygote present colored plumage, suggesting the W allele is recessive relative to the C allele.

**Fig. 5.**
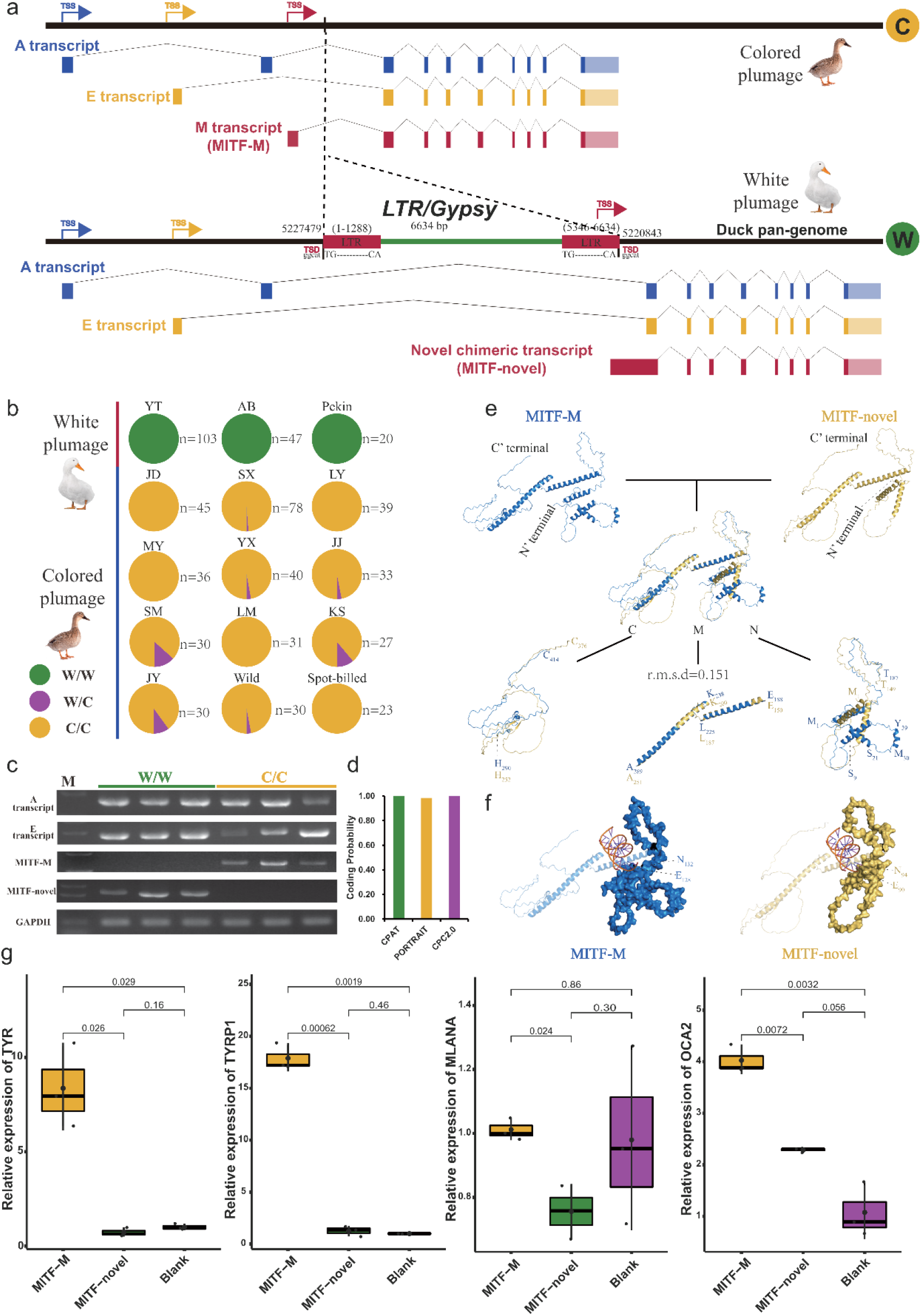
The novel chimeric transcript of *MITF* induced by a Gypsy insertion and its function. (a) The structure of the Gypsy that is anchored in the intron of *MITF* and identification of its transcript structure. (b) Genotypic frequencies of the *MITF* intron insertion in the validated populations with white and colored plumage by allelic-specific PCR genotyping. (c) Gel plots for the validation of identified transcripts using PCR amplification. (d) The Coding probability analysis for the MITF-novel transcript. (e) The overall structure and alignment of MITF-M and MITF-novel in cartoon mode. M represents the middle region of proteins, N represents the N-terminal region, and C represents the C-terminal region. (f) Amino acid binding characteristics of MITF-M and MITF-novel. The N-terminal regions show surface mode, and the rest show cartoon mode. The orange spiral structure indicates DNA fragments. (g) Quantitative PCR results showing the relative expression levels of four downstream genes of MITF in MITF-M over-expression cells MITF-Novel over-expression cells and blank.

The insertion of the W allele is predicted to be a member of the Gypsy superfamily, encoding two intact LTRs. However, intact proviral elements were not identified (Fig. 5a). This may be due to the accumulation of mutations following the TE insertion. Based on the sequence homology of the *LTR* region, this insertion occurred ∼0.6 million years ago, assuming the substitution rate of 1.91× 10^−9^ per site per year [40]. As intact proviral elements were absent, we investigated whether the intact LTR region might cause the effect of the W allele on *MITF*. We conducted RACE to identify the *MITF* transcripts in skin tissue from ducks at 1 day of age. In the SM duck with *MITF* CC genotype, three kinds of transcripts of *MITF* were identified, consisting of A, E and MITF-M (Fig. 5a, c). The MITF-M transcript of *MITF* is expressed exclusively in melanocyte lineages regulating the expressions of numerous pigmentation genes in the melanogenesis pathway and is responsible for melanocyte differentiation [41]. However, instead of A, E and MITF-M transcripts, we identified A, E, and a novel chimeric transcript (MITF-novel) in the skin tissue of YTG duck with WW genotype. This chimeric transcript fuses 151 bp 3’ end of the 3’ LTR, 878 bp 5’ flanking region of the exon 2, and 230 bp original exon 2. These three parts make up the novel exon 1. However, the MITF-M transcript was not found in the skin tissue of the YTG duck (Fig. 5a, c).

Coding probability analysis using CPAT, PORTRAIT, and CPC2.0 revealed that the MITF-novel transcript has a strong coding ability, supporting the existence of its protein product. Compared to the MITF-M transcript, MITF-novel lacks 39 amino acids at the N-terminal (Additional file 2: Fig. S7e). We predicted the tertiary structures to dissect the structural basis of the functional differences between MITF-M and MITF-novel (Fig. 5e-f). The structures of MITF-M and MITF-novel exhibit a substantially similar conformation in the middle region of the carbon backbone (E_188_-A_289_ of MITF-M in marine and E_150_-A_251_ of MITF-novel in yellow), with a root-mean-square deviation (R.M.S.D) value of 0.151. The structural differences were mainly concentrated in the loop regions at both N-terminal (N) and C-terminal (C) (Fig. 5e). The N-terminal 39 amino acids in MITF-M form two additional stable α-helix, S_9_-S_21_ and M_30_-Y_39_, around the nucleic acid binding region, according to the structure of 4ATI (Mus musculus) in the PDB database [42]. As can be seen from Fig. 5f, E90 and N94 in MITF-novel form a steric hindrance in conformation, thereby affecting the binding to the nucleic acid and might further block the downstream pathway. To further determine the regulatory role of the *MITF* gene in melanogenesis pathway, we transfected the MITF-novel and MITF-M transcript into DF-1 (chicken fibroblast cell line). Four well-known downstream genes of MITF-mediated melanogenesis pathway, including *TYR, TYRP1, MLANA* and *OCA2*, were significantly down-regulated in MITF-novel over-expressed cells relative to MITF-M over-expression cells (Fig. 5g). Except for the OCA2 gene, the expression level of the other three downstream genes in MITF-novel cells showed a similar pattern to the blank control (Blank). Additionally, these four downstream genes also had almost no expression in the skin tissue of Pekin duck (white plumage), but significantly higher expressions in Heiwu duck (colored plumage) (Additional file 2: Fig. S7f). This suggests that MITF-mediated melanogenesis genes may not be activated by MITF-novel, of which the lack of *TYR* expression might be the direct cause for the white plumage. The above evidence shows that the insertion of the Gypsy element induced the formation of MITF-novel transcript and the disruption of MITF-M, blocking the melanogenesis pathway and thus causing white plumage phenotype in WW genotype ducks.

## Discussion

Progress in characterizing genome-wide structural variations is shedding new light on how these mutations have impacted trait evolution in wild and domesticated species. To characterize duck genomic variation, we integrated five genome assemblies from wild and domesticated ducks to construct the first linear pan-genome using a combination approach of *Psvcp* [19] and *PPsPCP* [20] pipelines. This pan-genome can comprehensively capture genetic diversity by identifying a comprehensive set of SVs including those derived from TEs’ activity. Combining pan-genome, population-level sequencing data, and TE analyses enables us to resolve and link SVs to domestication and improvement traits. By identifying SVs associated with duck domestication and improvement, we demonstrated that genes adjacent to these SVs are enriched for growth, reproduction traits, and pathogen defense responses, which is in line with the intensive artificial selection in chicken [1]. The *IGF2BP1* loci demonstrate how this duck pan-genomic resource is essential to resolve complicated quantitative traits that were unsolved in previous SNP-based studies [35]. This finding reveals that the duck pan-genome provides an untapped resource of SVs associated with economically important traits, which can benefit QTL mapping and GWAS by capturing missing heritability and empowering breeding.

Multiple studies revealed that SVs had pronounced phenotypic impacts via disrupting gene function or regulating gene expression dosage [43]. According to our pan-genome analysis, TEs are an important source for the generation of SV. These can have large phenotypic impacts due to their innate structure that comprises proviral elements capable of donating promoters or enhancers that regulate transcriptional networks or generate a novel transcript [9]. A Gypsy element inserted in the *IGF2BP1* promoter provide an agonist to increase its transcript, leading to enlarging bodyweight. A Gypsy element anchored in the intron of *MITF* generated a novel transcript, leading to melanin absence and white plumage formation. These two examples of TE-derived SVs with major phenotypic impacts illustrate these mutations can shape both quantitative and qualitative traits. Similar findings were also reported in other species. For example, a recent study on Norwegian sheep found that a TE insertion could suppress the beta-carotene oxygenase 2 (*BCO2*) gene expression and result in yellow adipose tissue [44]. A LINE insertion can increase the transcriptional activity of the agouti signaling protein (*ASIP*) gene responsible for the white coat color in swamp buffalo [12]. A TE insertion in the intron 4 of Tyrosinase (*TYR*) gene also induced aberrant transcripts leading to a recessive white plumage in chickens [45]. In plants, a pan-genome study of tomatoes revealed hundreds of SVs, most of which were TE-related, linked to expression changes in associated genes, with potentially large impacts on quantitative trait variation [46]. Combining these examples and findings in our study, we suggest that TE-derived SVs have a pronounced phenotypic impact on quantitative and qualitative traits, affecting trait formation in livestock. Since TEs exist in almost all eukaryotic genomes and carry pronounced phenotypic impact, TE-derived SVs can be used as additional markers to facilitate causal variant identification for target phenotype and further genetic improvement through genomic selection and marker-assisted selection.

## Conclusions

In summary, we constructed the first duck pan-genome and further identified a comprehensive set of SVs in duck. A substantial number of SVs were derived from TE activity, which was linked to domestication and improvement traits. To validate the pronounced phenotypic effect of TE-derived SV during evolution, we dissected one TE-derived SV located at promoter region of *IGF2BP1* enlarging bodyweight by increasing its expression, and revealed another TE-derived SV located within intron of *MITF* responsible for white plumage formation via generating a novel transcript. Our findings highlight the importance of TE-derived SVs’ pronounced impact on both quantitative and qualitative traits and will shed the light on how TE-derived SVs impact phenotype formation in animals.

## Materials and Methods

### Duck genome and genomic sequencing

The reference duck genome (ZJU1.0) [47] and four chromosome-level duck genomes were downloaded from the National Center for Biotechnology Information (NCBI) and ENSEMBL database, which consists of the duckbase.refseq.v4, CAU_Pekin_2.0, CAU_Laying_1.0 and ASM874695v1 genomes [48] (Additional file 1: Table S1). This study generated in-depth genomic sequencing data of 131 unrelated duck individuals, including two wild breeds, seven Chinese indigenous breeds and three commercial breeds (Additional file 1: Table S2). Genomic DNA was extracted from duck blood samples using Qiagen DNeasy Kit. Paired-end libraries with ∼500 bp insertion size were constructed and then subjected to sequencing using the BGISEQ-500 platform to generate paired-end 150 bp reads (BGI Genomics Co., Ltd., China).

### Pan-genome construction

The relationships of colinear chromosomes between the four duck genomes and the reference genome were identified using the MCscan (https://github.com/tanghaibao/jcvi/wiki/MCscan-(Python-version)). We first used the *Psvcp* pipeline [19] to construct the duck Pan-genome.1 (Fig. 1a). Briefly, query chromosome was aligned on the reference chromosome using nucmer command in mummer (v4.0.0beta2) [49] (Additional file 2: Fig. S1b), then subjected to structural variants detection using Assemblytics [50]. The insertions with a size of more than 50 bp were placed into the reference chromosome. The annotation file of reference was also updated, along with the integration of insertions. The four query duck genomes duckbase.refseq.v4, CAU_Pekin_2.0, CAU_Laying_1.0 and ASM874695v1 were aligned on the reference ZJU1.0 one by one. Secondly, the four query genomes were also aligned on the Pan-genome.1 to identify the novel sequence using the *PPsPCP* pipeline [20].

All novel contigs were merged and then redundant assembled sequences were filtered using CD-HIT [51] (-c 0.9 -aS 0.8 -d 0 -sf 1) with the threshold of 90% similarity. Newly contigs of non-reference sequences with lengths larger than 500 bp were kept. Novel contigs were aligned using blastn (v2.9.0) [52] against the NT database (v5, 07-03-2019) of contaminant taxid groups, which includes archaea, viruses, bacteria, fungi and Viridiplantae to identify the contaminant sequences. However, we could not find any hits with identities larger than 90% and query lengths larger than 50%. The final contamination-free non-reference sequences and the Pan-genome.1 were merged to generate the duck pan-genome. The novel contigs were annotated with ASM874695v1 annotation file using the GMAP(v2021-08-25) [53].

### PAV calling and pan-genome modeling

Steps for Presence/Absence Variation (PAV) calling was described in our previous study [1]. Briefly, the longest transcripts of each gene were retrieved, and coding sequence (CDS) regions were extracted. Genes that cumulative coverage of at least two reads with more than 5% of all exons was considered as presence, otherwise absence [54]. Clean reads were aligned to the duck pan-genome using BWA-MEM (v0.7.17) [55] and the sequencing depth was counted using Mosdepth package (v0.2.5) [56]. It is confirmed that the sequencing data with more than 10x in each depth was allowed to obtain a PAV matrix with a 99.4% accuracy rate [1]. In this study, the average depth of all sequencing data was more than 45x, which increased the robustness of PAV calling. Pan-genome modeling was performed by curve fitting of the pan-genome and core genome. A pan-genome curve was constructed using a power-law regression: y = AxB + C. A core genome curve was performed using an exponential regression model: y = AeBx + C. In these two equations, y was the total number of the gene, x was the genome number, and A, B, and C were the fitting parameters.

### SV calling

To facilitate the accurate alignment of sequencing reads at the boundaries of novel contigs, 200 bp flanking sequences on either side were added [57]. High-depth sequencing data was mapped on the duck pan-genome with flanking sequencing. Bam files were sorted, and duplicate reads were removed using Sambamba (v0.8.2) [58].

Structural variants were called using four commonly used SV detecting tools relying on different bioinformatics algorisms, including LUMPY (v0.2.13)[23], Delly (v0.8.7)[24], GRIDSS (v 2.1.0) [25] and Manta (v1.6.0)[26]. LUMPY generated deletions, which were less than 340 bp and no split read support, were filtered to reduce the false calls [28]. Subsequently, the VCF files were genotyped using SVTyper (v0.0.4)[59]. We used Delly, GRIDSS, and Manta to identify and genotype the SVs with default parameters. SV genotypes from four tools were merged and filtered using SURVIVOR [60] with the parameters “ SURVIVOR merge 1000 2 1 1 0 50 “ to keep the variants detected by at least two tools.

### SV analysis

Population genetic analysis was conducted using the binary SV genotype. Biallelic SVs located on autosomes were retained and subjected to filter against the variants with MAF< 0.01 using PLINK (v1.9) [61]. A phylogenetic tree was constructed using the IQ-TREE software (v1.6.12) [62] with 1000 bootstrap replicates based on the GTR model and visualized using the iTOL online web server [63]. PCA was implemented using smtpca of the EIGENSOFT [64]. Population assignment analysis was conducted using the Admixture software [65].

To identify the SVs with significantly changed occurrence frequency during domestication or improvement, the derived allele frequencies were compared between the native breeds and wild breeds or commercial breeds. The wild group comprised Mallard duck and Chinese-spot-billed duck, while the commercial group consisted of Pekin duck, Cherry Valley duck, and Grimaud freres duck. The other seven populations were indigenous duck breeds, defined as the native group. The significance of the difference in frequencies for each SV between groups was determined by Fisher’s exact test with a false discovery rate (FDR) of 0.001 [1, 30]. Significantly increased SVs during domestication or improvement, were defined as SVs having a significantly higher frequency in native breeds than wild breeds, or commercial breeds than native breeds, respectively. Inversely, we consider SVs with a significantly lower frequency as significantly decreased SVs.

### Detection of transposable elements detection and their association with SVs

TEs can be classified into two classes: DNA transposons and retrotransposons, depending on genetic structures and transposition mechanisms. Long terminal repeats (LTRs), long interspersed nuclear elements (LINEs), and short interspersed nuclear elements (SINEs) are the dominant retrotransposons in vertebrates [10]. Interspersed repeats and low complexity DNA sequences across the duck pan-genome were screened using RepeatMasker (v4.0.8) [66]. A custom repeat library was constructed using RepeatModeler (v1.0.11) [67]. Repeat sequences detected based on the custom repeat library and one the RepBase database (downloaded in June 2019) of vertebrates were scanned separately and merged using ProcessRepeats program of RepeatMasker. Abundance, length, and divergence rate were abstracted from the result file of RepeatMasker (v4.0.8) [66]. TE families with copy numbers more than 2,000 were collected to construct the classification tree. The top 17 class/superfamilies filtered according to their copy numbers were retrieved to perform the statistics of length, abundance, and divergence levels. To examine the occurrence correlation between SVs and TEs, we divided the genome into windows with 2 kb, 5 kb and 10 kb and subject to screen the presence or absence of TE and SV. The occurrence matrix was subjected to the Chi-square test using the R package ‘ggstatsplot’ [68].

### Identification of intact TEs and estimation of insertion time

Intact TEs were detected using Extensive De-novo TE Annotator (EDTA) [69] with default parameter. Structural and proviral elements were resolved by RetroTector [70] and EDTA software. Insertion time of LTR TEs was estimated with the model of T = K/(2r) (https://github.com/wangziwei08/LTR-insertion-time-estimation), assuming that the substitution rate is 1.91 × 10^−9^ per site per year [40].

### GO annotation

To identify the genes affected by SVs, bedtools (v2.29.2)[71] was used to search for genes physically closest to SVs. Functional annotation of the duck pan-genome was performed using the command line Blast2GO (v2.5)[72]. The longest transcript of each gene was retrieved from the pan-genome and subjected to alignment to the proteins in the Uniref90 database (downloaded on Mar 2022) using BLASTP function in Diamond [73] with the threshold E-values < 1×10^−3^. Gene ontology annotation of these genes was conducted by the R package topGO [74] using Fisher’s exact test with the approach ‘elim’ for multiple comparisons correction.

### Calculation of absolute allelic frequency differences (absAFdif)

Of the WGS data from 131 duck individuals used in this study, 41 are from commercial meat breeds (Pekin, Cherry Valley, and Grimaud freres ducks) with high body weight and thus categorized into the high group, and other 90 individuals were categorized into the low group. Variants located within the 10 kb upstream and 10 kb downstream regions (pan-genome chr28:103034-130138) of the *Gypsy* element were extracted. The absAFdif value for each variation (SNPs and SV) was calculated by the comparison of allele frequencies between the two groups.

### Genotyping of SVs and association analysis

Primers were designed to genotype the SVs based on the flanking sequence (Additional file 1: Table S8). PCR genotyping was described as our previous study [1]. A general linear model (GLM) was conducted to investigate the association between *IGF2BP1* genotypes and phenotypes using TASSEL5 [75] software with sex factor defined as a fixed effect. The value of marker R-squared was determined to explain the phenotypic variation derived from genotypes, which was computed from the marker sum of squares after fitting all other model terms divided by the total sum of squares. A total of 8 trait phenotypes of Liancheng white duck × Kaiya duck F2 population were collected [76].

They consist of 3 bodyweight traits including bodyweight at ages of the week of 8 and 9 (BW8 and BW9) and BWHR (bodyweight of feather removed at slaughter), and 5 carcass composition traits including evisceration weight (EW and SEW), leg weight (LW), breast muscle weight (BMW) and carcass weight (CW).

### Functional assay of *IGF2BP1* and *MITF*

Rapid amplification of cDNA ends (RACE) was conducted to amplify the full length of *IGF2BP1* and *MITF* transcripts using SMARTer Race 5’/3’ Kit with gene specific primers (Additional file 1: Table S8). To further verify the molecular effects of the insertion, luciferase expression levels were investigated to represent the transcriptional activity through transfecting two kinds of recombinant plasmids (PGL3-L and PGL3-L+LTR). A recombinant plasmid pGL3-L+LTR was defined to represent the TE insertion as its incapacity to insert the whole TE into the pGL3 plasmid. Promoter region of *IGF2BP1* was cloned into the PGL3-Basic luciferase vector (Promega) that was subjected to transfection into DF-1 cell line (chicken fibroblast cell) together with PRL-TK plasmid. Transcriptional activity was investigated by Dual-Luciferase Reporter Assay System (Promega) after 48 hours of transcription. The mRNA level of *IGF2BP1* and *MITF* were determined by quantitative PCR with their specific premiers (Additional file 1: Table S8), normalized by *GAPDH* gene using the 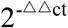 method. AlphaFold2 was used to predict different conformational structures for MITF-M and MITF-novel [77, 78]. Protein structural analysis and all colored schemes are accomplished by PyMOL 2.5. The 3xFLAG sequence was inserted into the N-terminal of the MITF-M and MITF-novel transcripts and then transfected into the DF-1 cell line. After 48 hours of transfection, their regulatory effects on the four well-known downstream genes in the MITF-mediated melanogenesis pathway were determined using the quantitative PCR (Additional file 1: Table **S8**).

### RNA-Seq analysis

Transcriptome data of liver and sebum from mallard and pekin at 2, 4, and 6 weeks of age was downloaded from National Genomics Data Center (China National Center for Bioinformation) from the project number PRJCA002795, PRJCA002803, PRJCA002808, and PRJCA002807 [48]. Transcriptome data of the skin tissue was downloaded from project number PRJCA003516 [79]. Raw reads were filtered using Trimmomatic (v 0.39) [80] and subjected to align the duck pan-genome using STAR (v2.7.1a) [81]. Expression levels of transcripts were determined using featureCounts (v2.0.0) [82].

## Supporting Information

**Additional file 1: Supplementary Table S1-S8**.

**Additional file 2: Supplementary Figures S1-S7**.

**Additional file 3: Supplementary Notes**.

## Acknowledgements

We thank Longxian Zhang, Jianying Huang and Jiang Li for helping with computing resources. We also thank National Supercomputing Center in Zhengzhou and High-Performance Computing platform in College of Veterinary Medicine of Henan Agricultural University for providing computing resources.

## Authors’ contribution

W. L., K. W., X. K. and H. H. conceived the Project and designed research. K.W., and W. L. designed the analysis and wrote the manuscript; K. W., W.L., H. H., J. L., and A.S. the revised manuscript. K. W., G. H., J. L. and Y. W. performed analysis; W. L., C. Z., Y. L., and X. H. performed the wet-lab experiment; Y. Y., P. G., K. W., Y. G., S. Z., Y. F., T. Z., L. L., X.K., Y.T., G.S., and R. J. contributed to sample collection and construction of F2 resource population.

## Availability of data and materials

All the sequence data generated in this study have been deposited in the National Genomics Data center (https://bigd.big.ac.cn) with the accession codes PRJCA011446. The duck pan-genome and relevant data are available in the DRYAD database (https://doi.org/10.5061).

## Ethics declarations

Ethics approval for this study was obtained from Henan Agricultural University.

## Consent for publication

Not applicable.

